# Exome chip meta-analysis elucidates the genetic architecture of rare coding variants in smoking and drinking behavior

**DOI:** 10.1101/187658

**Authors:** Dajiang J. Liu, David M. Brazel, Valérie Turcot, Xiaowei Zhan, Jian Gong, Daniel R. Barnes, Sarah Bertelsen, Yi-Ling Chou, A. Mesut Erzurumluoglu, Jessica D. Faul, Jeff Haessler, Anke R. Hammerschlag, Chris Hsu, Manav Kapoor, Dongbing Lai, Nhung Le, Christiaan A de Leeuw, Ana Loukola, Massimo Mangino, Carl A. Melbourne, Giorgio Pistis, Beenish Qaiser, Rebecca Rohde, Yaming Shao, Heather Stringham, Leah Wetherill, Wei Zhao, Arpana Agrawal, Laura Beirut, Chu Chen, Charles B. Eaton, Alison Goate, Christopher Haiman, Andrew Heath, William G. Iacono, Nicholas G. Martin, Tinca J. Polderman, CHD Exome+ Consortium, Consortium for Genetics of Smoking Behavior, Alex Reiner, John Rice, David Schlessinger, H. Steven Scholte, Jennifer A. Smith, Jean-Claude Tardif, Hilary A. Tindle, Andreis R van der Leij, Michael Boehnke, Jenny Chang-Claude, Francesco Cucca, Sean P. David, Tatiana Foroud, Sharon L.R. Kardia, Charles Kooperberg, Markku Laakso, Guillaume Lettre, Pamela Madden, Matt McGue, Kari North, Danielle Posthuma, Timothy Spector, Daniel Stram, David R. Weir, Jaakko Kaprio, Gonçalo R. Abecasis, Scott Vrieze

**Author notes:** These authors contributed equally to the work. Address correspondence to Scott Vrieze, Department of Psychology, University of Minnesota. 75 East River Road, Minneapolis, MN 55455. See supplement for a list of authors associated with the replication consortia.

## Abstract

**Background:** Smoking and alcohol use behaviors in humans have been associated with common genetic variants within multiple genomic loci. Investigation of rare variation within these loci holds promise for identifying causal variants impacting biological mechanisms in the etiology of disordered behavior. Microarrays have been designed to genotype rare nonsynonymous and putative loss of function variants. Such variants are expected to have greater deleterious consequences on gene function than other variants, and significantly contribute to disease risk.

**Methods:** In the present study, we analyzed ∼250,000 rare variants from 17 independent studies. Each variant was tested for association with five addiction-related phenotypes: cigarettes per day, pack years, smoking initiation, age of smoking initiation, and alcoholic drinks per week. We conducted single variant tests of all variants, and gene-based burden tests of nonsynonymous or putative loss of function variants with minor allele frequency less than 1%.

**Results:** Meta-analytic sample sizes ranged from 70,847 to 164,142 individuals, depending on the phenotype. Known loci tagged by common variants replicated, but there was no robust evidence for individually associated rare variants, either in gene based or single variant tests. Using a modified method-of-moment approach, we found that all low frequency coding variants, in aggregate, contributed 1.7% to 3.6% of the phenotypic variation for the five traits (p<.05).

**Conclusions:** The findings indicate that rare coding variants contribute to phenotypic variation, but that much larger samples and/or denser genotyping of rare variants will be required to successfully identify associations with these phenotypes, whether individual variants or gene‐based associations.

## Introduction

Tobacco and alcohol use together account for more morbidity and mortality in Western cultures than any other single risk factor or health outcome^1^. These preventable and modifiable behaviors are heritable^2^, have been associated previously in human and model organism research with multiple common genetic variants^3-7^, and most prominently feature genes involved in alcohol/nicotine metabolism and nicotinic receptors.

Advances in sequencing technology have led to cost-effective “exome arrays”, which affordably genotype a few hundred thousand rare (minor allele frequency [MAF]<1%), putatively functional exonic variants. Compared to common SNPs (MAF>1%) used in genome-wide association studies (GWAS), rare exonic variants may have greater potential to elucidate the biological mechanisms of addiction and other complex traits^8^, ^9^. Loss of function (LoF) variants result in the loss of normal function of a protein, and may have greater phenotypic impact than other variants that do not have obvious biological consequences^10^, ^11^. One well-known example is rare LoF mutations in *PCSK9* that greatly reduce risk of cardiovascular disease with no apparent negative effects, encouraging the development of a new class of *PCSK9* inhibitor drugs^12^, ^13^.

The analysis of any rare event, including rare genetic variants, presents analytical challenges. First, statistical power is a function of MAF, such that rare variants of small to moderate effect require very large samples to achieve adequate statistical power^14^. Statistical association techniques have been developed to mitigate this issue, including tests that aggregate information across many low-frequency variants^15^. These “burden” tests can improve power under certain assumptions, such as that a large proportion of the aggregated variants are independently associated with the phenotype of interest. Here, we use novel methods to implement a variety of genetic association tests, in the largest sample currently available, to test the effect of rare and low-frequency exonic variants on tobacco and alcohol use behaviors.

The vast majority of existing addiction-related rare variant studies use targeted sequencing of known addiction-associated loci to discover and test for association. This has led to intriguing new leads, especially within nicotinic receptor gene clusters^16-25^ and alcohol metabolism genes^26-28^ for alcohol and nicotine dependence. This strategy has also produced rare variant associations with alcohol dependence in genes not previously implicated in addiction. In one case, burden testing was used to find an association with rare variants in *SERINC2*^28^. In another case, a burden test across *PTP4A1*, *PHF3*, and *EYS* showed an association with alcohol dependence^29^. Single variant tests did not reach significance after multiple-testing corrections in either case. In part due to the nature of burden tests, especially when conducted across multiple genes, these findings do not have simple biological interpretations, and no rare variant results have been replicated.

Some studies also leverage information about predicted functional consequences of rare mutations to increase the power of association analyses. For example, one study of nicotine dependence found significant rare single-variant associations in *CHRNB4*, but only when variants were weighted by their effect on the cellular response to nicotine and acetylcholine^30^. Such positive findings benefit from replication, which has not always been straightforward. For example, all rare variant associations in addiction are, to our knowledge, candidate gene analyses with type I error thresholds based only on tests within that region. Historically, such analyses have tended to produce overly optimistic estimates of the number of associated loci^31^. Genome-wide analyses with more conservative type I error thresholds have reported null rare variant findings across an array of phenotypes relevant to addiction^32-34^. Precisely because genome-wide analyses are conducted on many variants across the genome, they are in principle able to discover novel rare variant associations within new or known loci. One way to improve power in genome-wide analyses is through genetic association meta-analysis, which entails the aggregation of results across many studies to achieve large sample sizes. We present here such a meta-analysis, aggregating studies with rare variant genotype arrays and measured alcohol and nicotine use, to arrive at a highly powered test of the hypothesis that rare exonic variants affect addiction-related outcomes.

In addition to single variant and gene-level tests, we also conducted tests of the contribution of rare nonsynonymous variants to the heritability of our alcohol and tobacco use phenotypes. Twin studies, as well as studies of the aggregate effects of common variants, have found both alcohol use and tobacco use to be heritable behaviors^34-39^. Research on the aggregate contribution of rare variants, however, has been scarce, with previous work on related phenotypes in smaller samples failing to detect aggregate effects for smoking and alcohol consumption^32^. In this study, we implemented a novel method-of-moments approach to analyze heritability and genetic correlations due to variants genotyped on the exome array. We used meta-analytic summary statistics to quantify the contribution to heritability of variants in various functional categories and frequency bins, estimated the genetic correlation between smoking and drinking traits, and evaluated the contribution of rare coding variants to the phenotypic variation of smoking and alcohol use behavior.

## Methods and Materials

Seventeen studies contributed summary statistics for meta-analysis. These studies, their sample sizes, and available phenotypes are listed in the online supplement (Tables S1 and S2). Two studies (HRS and COGA) provided results for individuals of European and African ancestry separately. One study (the UK Biobank of European ancestry) was stratified into two samples according to ascertainment protocol and genotyping method. Thus, in the end, 20 independent sets of results from 17 independent studies were submitted for meta-analysis.

### Ancestry

Allanalyseswerestratifiedbyancestry.EighteendatasetswereonindividualsofEuropean ancestry and two datasets on individuals of African ancestry.

### Phenotypes

Phenotypes were selected to be relevant to prior GWAS of smoking and alcohol use, common in psychological, medical, and epidemiological data sets, and known to be correlated with measures of substance dependence. Five phenotypes were selected based on their inclusion in previous successful GWAS studies^4^, ^40-42^ and availability among large exome chip studies.

1. *<u>Cigarettes per day</u>*. The average number of cigarettes smoked in a day among current and former smokers. Studies with binned responses retained their existing bins. Studies that recorded an integer value binned responses into one of four categories: 1=1-10, 2=11-20, 3=21-30, 4=31 or more. Anyone reporting 0 cigarettes per day was coded as missing. This phenotype is a component of commonly used measures of nicotine dependence such as the Fagerstrom Test for Nicotine Dependence.
2. <u>*Pack Years*</u>. Defined in the same way as cigarettes per day but not binned, divided by 20 (cigarettes in a pack), and multiplied by number of years smoking. This yields a measure of total overall exposure to tobacco and is relevant to cancer and chronic obstructive pulmonary disease risk.
3. <u>*Age of Initiation of Smoking*</u>. A measure of early cigarette use. Defined as the age at which a participant first started smoking regularly.
4. <u>*Smoking Initiation*</u>. A binary variable of whether the individual had ever been a regular smoker (1) or not (0), and often defined as having smoked at least 100 cigarettes during one’s lifetime.
5. <u>*Drinks per week.*</u> A measure of drinking frequency/quantity. The average number of drinks per week in current or former drinkers.

### Genotypes

Fifteen of the seventeen studies were genotyped with the Illumina HumanExome BeadChip, which contains ∼250,000 low-frequency nonsynonymous variants, variants from the GWAS catalog, and a small number of variants selected for other purposes. Two studies were genotyped on the Illumina Human Core Exome, which includes an additional ∼250,000 tag SNPs. Finally, the present study used the initial release of 150,000 UK Biobank participants, which comprised two cohorts: 1) the UK BiLEVE cohort of ∼50,000 heavy and never smokers genotyped on the UK BiLEVE array and 2) ∼100,000 participants genotyped on the UK Axiom array. These arrays are highly similar and described elsewhere (http://www.ukbiobank.ac.uk/scientists-3/uk-biobank-axiom-array). They both include >800,000 variants including ∼630,000 genome-wide tagging markers, ∼110,000 rare coding variants that are largely a subset of variants also genotyped on the Illumina HumanExome Beadchip, and an additional ∼107,000 variants chosen for the study of specific medical conditions.

### Generation of Summary Association Statistics

Twenty sets of results from 17 independent studies (Table S1) with smoking and drinking phenotypes were included in the discovery phase. Summary statistics were adjusted by local analysts for age, sex, any study-specific covariates, and ancestry principal components (see Table S2 for genomic controls). For studies with related individuals (see Table S1), relatedness was accounted for in linear mixed models using empirically estimated kinships from common SNPs^43^. Residuals were inverse-normalized to help ensure well-behaved test statistics for rare variant tests.

Quality control of per-study summary statistics included evaluation and correction of strand flips and allele flips through systematic comparison of alleles and allele frequencies against reference datasets ExAC v2.0, 1000 Genomes Phase 3, and dbSNP. Variants with call rates<0.9, and Hardy Weinberg *p*<1x10^−7^, and polymorphic in <3 studies, were also removed. The latter filter was meant to avoid findings that could not be broadly replicated across the 17 studies.

Variants were annotated against RefSeq 1.9^44^. The allelic spectrum of all nonsynonymous, start loss/gain, stop loss/gain, or splice acceptor/donor is displayed in Figure 1 for cigarettes per day, stratified by whether the variant exists only in the UK Biobank, only in studies genotyped with the Illumina exome chip, or in both. More details on the allelic spectra within functional classes are available in Table S3 and Figure S1.

**Figure 1.**
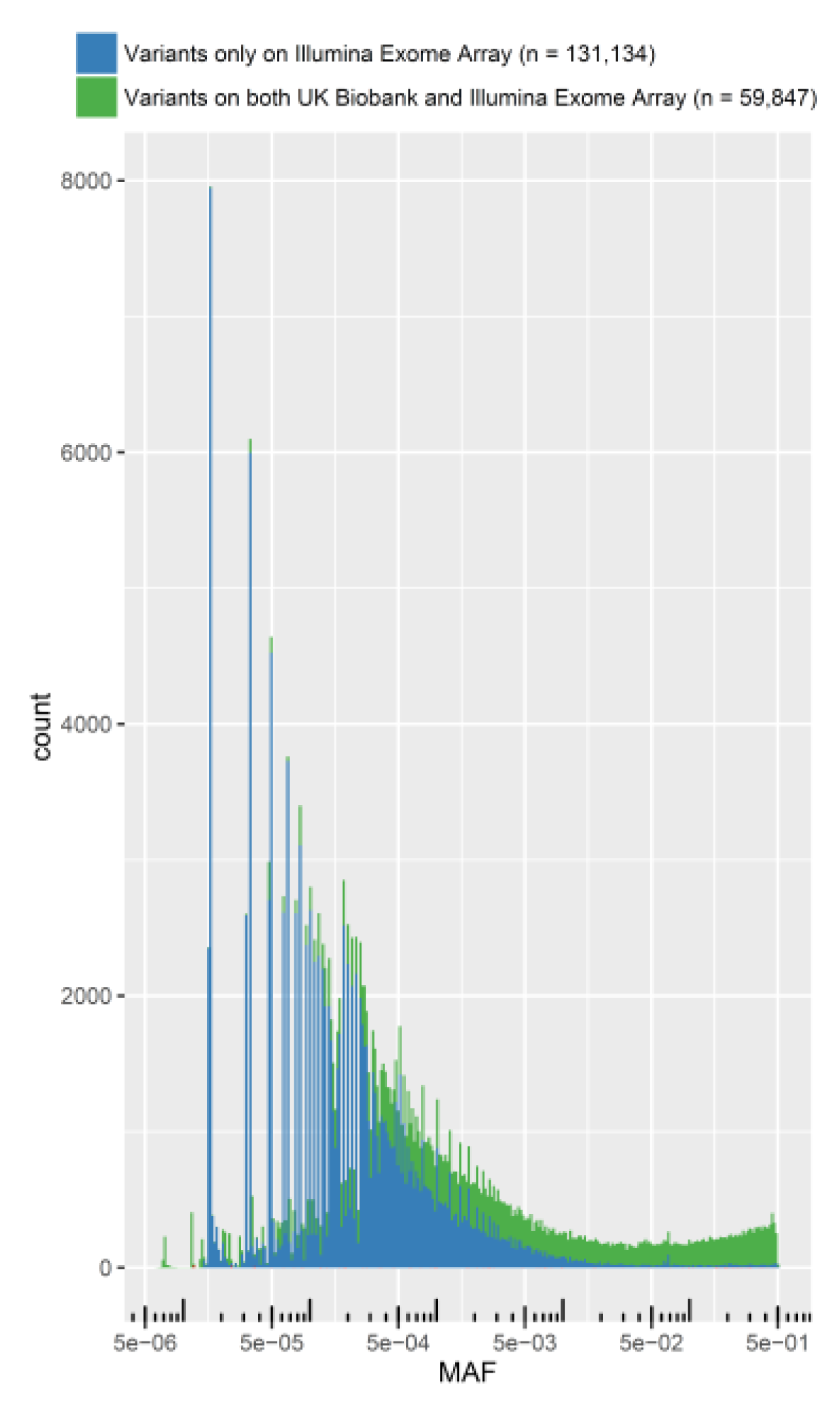
Distribution of nonsynonymous and loss of function variant allele frequencies in the Illumina exome array and the UK Biobank arrays, generated from the results for cigarettes per day. (Allelic spectra for other phenotypes may differ slightly). Note there are only 241 variants that were present only in the UK Biobank and not on the Illumina Exome Chip; these 241 variants are not displayed in the figure. MAF = minor allele frequency estimated in the meta-analysis.

### Meta-analysis

We performed meta-analysis in rareMETALS version 5.8^45^ using the Mantel-Haenszel method^45^. For gene-level burden tests, we selected variants predicted to be nonsynonymous, start loss, start gain, stop loss, stop gain, or splice donor/acceptor within each gene from RefSeq 1.9^44^. Two complementary gene-level association tests were performed: the sequence kernel association test (SKAT; ^46^, ^47^) with MAF cutoff 1% and a variable MAF threshold test (VTCMC; ^48^) with a maximum MAF=1%. We chose variants with MAF≤1% as we were interested in the contribution of variants with a frequency lower than that which has been reliably imputed and tested in past GWAS meta-analyses. Exceedingly rare variants, with minor allele counts less than five, were excluded from single variant analyses due to extremely low expected power. These rare variants were included in all gene-based tests.

There exist known genetic associations between common variants and smoking or drinking phenotypes, including variants within the nicotinic receptor gene cluster on chromosome 15 with cigarettes per day^4^ ^5^; *CYP2A6* and *CYP2B6* with cigarettes per day^42^; *AUTS2, KLB, ADH1B, ALDH2,* and *GCKR* with alcohol use^40^ ^41^; and *NCAM1* and *TEX41* with smoking initiation^49^. We conducted sequential forward selection association tests, as implemented in rareMETALS, for rare variants within these regions, controlling for any common variant associations in these regions.

Association testing was done in stages. First, we tested common variants within all known loci associated with these phenotypes, as listed above. To these variants we applied the standard genome-wide significance threshold of *p*<5e-^8^. Second, for rare variants with MAF<1% and minor allele count ≥5, we applied a Bonferroni correction for the number of such variants tested, resulting in p-value thresholds from 2.1x10-^7^ to 2.2x10-^7^ depending on the phenotype. This threshold was applied to both marginal (unconditional) analyses and forward selection conditional analyses of rare variants within known loci.

Third and finally, for each known and previously validated locus associated with these and related traits, we explored a relaxed multiple-testing threshold based only on the number of rare variants within a 1MB region around the most highly significant (usually common variant) association within that region. Each locus-wide p-value threshold is provided in Table 1. This approach is meant to mimic the typical candidate gene or targeted sequencing approach, where one or a few known loci are analyzed separately from the rest of the genome. While this threshold is overly liberal, it allows a more direct comparison between our results and existing publications of rare variants described in the introduction.

**Table 1.**
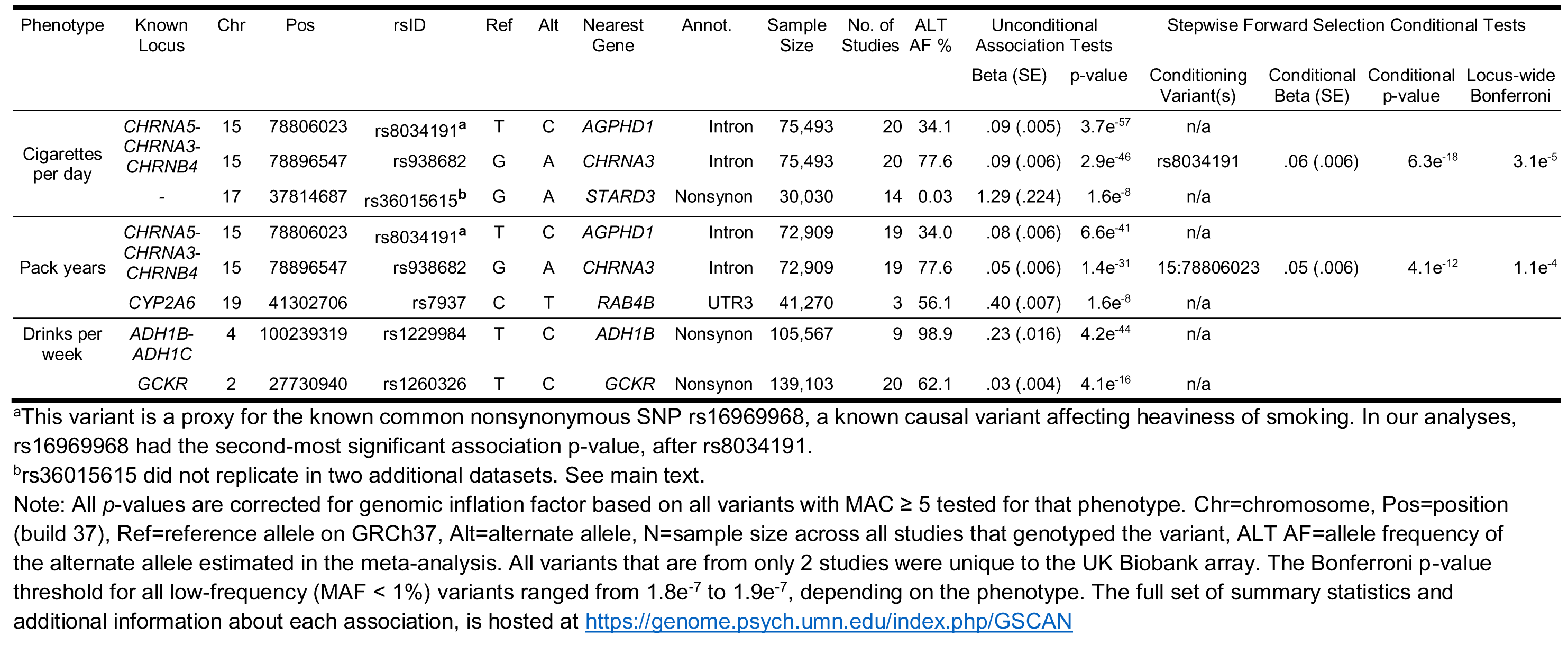
TR results for common and rare (MAF < 1%) variants. Unconditional results and conditional results based on stepwise forward selection are shown.

Finally, we attempted to replicate previous rare variant associations referenced in the introduction and listed in Table S6. The prior studies were of alcohol or nicotine dependence. We attempted replication in our phenotypes for any single variant when that variant was included on the exome array (5 of 23 variants were available) and, if not, we took the variant with the smallest p-value within the same gene as the original finding (16 of remaining 18 variants), or any gene-based burden test for which content existed on the array (27 of 27 prior associated genes had content on the array). We applied a liberal threshold that corrected only for the number of tests conducted for this replication exercise (.05/52=.00096).

### Replication Data

We replicated any novel exome-wide significant rare variant (MAF<1%) in two additional exome chip smoking meta-analysis efforts, the CHD Exome+ Consortium (N=17,789) and the Consortium for Genetics of Smoking Behaviour (N=28,583). Both consortia defined their phenotypes similarly and corrected for sex, age, principal components (and/or genetic relatedness, as appropriate), and inverse-normalized prior to association analysis.

### Genetic Architecture Analysis

We performed heritability and genetic correlation analyses using a modified method-of-moment estimator, adapated to the analysis of sparsely genotyped rare variants. The method calculates covariate-adjusted LD scores from summary statistics based upon partial correlations and quantifies the uncertainty of LD scores with a bootstrap procedure that uses multiple contributing studies. The estimation of heritability follows established methods. Detailed descriptions of the approach can be found in the supplement.

## Results

Conditionally independent, significant single-variant association results are displayed in Table 1. We discovered a single novel association signal for a single rare variant, only for cigarettes per day, rs36015615 (N=30,030; Beta=1.3; *p*=9.5x10^−9^), a nonsynonymous SNP in the gene *STARD3*. This novel variant did not replicate in either of two replication consortium datasets, the CHD Exome+ Consortium (N=17,789, Beta=-.01, *p*=.94) or the Consortium for Genetics of Smoking Behaviour (N=28,583, Beta=.056, *p*=.84).

Two known common variants within the *CHRNA5-CHRNA3-CHRNB4* locus (rs16969968 and rs938682) were independently associated with cigarettes per day and pack years. Conditional tests of rare nonsynonymous variants within these genes were non-significant. We verified at *p*<5e^−8^ a common variant association near *CYP2A6* for pack years. For drinks per week, we replicated at *p*<5e^−8^ a variant in *GCKR* and a known low-frequency association for a nonsynonymous variant in *AHD1B*, but did not replicate at *p*<5e^−8^ prior genome-wide associations around *AUTS2*. In *GCKR* we discovered a common nonsynonymous SNP, rs1260326, associated with drinks per week. This SNP is 10,047 base pairs from, and in high LD (r^2^=.97 in 1000 Genomes Phase 3 data) with the intronic *GCKR* SNP rs780094 that almost reached statistical significance (*p*=1.6x10^−7^) in a recent report^40^.

We removed variants that were only present in two or fewer studies to avoid reporting associations that arose solely from the UK Biobank and are essentially unreplicable. This filter removed several genome-wide significantly associated common variants previously reported to associate with either tobacco or alcohol use. These variants included rs1137115 in *CYP2A6* associated with cigarettes per day as well as some rare variants within that locus that showed evidence of conditionally independent association. Additional UK Biobank-associated variants were rs4144892 in *NCAM1* associated with SI; rs58930260, rs11694518, and rs12619517 associated with SI; rs12648443 in *ADH1C* associated with drinks per week; and rs13146907 in an intron of *KLB* associated with drinks per week. These results are reported in Table S4 of the supplement.

SKAT gene-based tests of nonsynonymous variants with MAF<1% resulted in one significant association with *ADH1C* (SKAT *p*=1.0x10^−8^), although after conditioning this gene-based test on a nearby genome-wide significant nonsynonymous variant in *ADH1B*, the *ADH1C* effect becomes nonsignificant (*p*=.52). Variable threshold gene based burden tests (VTCMC) yielded no significantly associated gene.

Out of four published rare single variant associations with addiction phenotypes, we replicated only one, even after examining other variants in the same gene and applying a relaxed multiple testing threshold (Table S6). The variant was rs56175056 in *CHRNA4* (*p*=9.4e^−6^), previously identified in an Icelandic population^50^. Out of twenty six genes that have been associated with alcohol or nicotine dependence in published rare variant burden tests, we found a significant association for one, *ADH1C* (Table S6), described in the previous paragraph.

Heritability was estimated for each trait and partitioned by annotation category. First, we annotated variants on the exome chip based upon gene definitions in RefSeq 1.9, using SEQMINER version 6.0^51^. Sixteen functional categories were considered, including downstream, essential splice site, noncoding exon, intergenic, intron, common nonsynonymous (MAF>0.01), rare nonsynonymous (MAF<0.01), normal splice site, start gain/loss, stop gain/loss, synonymous, and 3’/5’ untranslated regions. We fitted the baseline model with 16 categories, and estimated phenotypic variance explained by each category (Table S5).

Significant phenotypic variance was explained from rare nonsynonymous variants for all traits (p<0.05), from 1.7%-3.6% (Table 2). We also estimated the phenotypic variance explained by all variants on the exome chip, through aggregating the variance explained by each significant category (p<0.05) listed in Table S5. The total variance explained was highest for cigarettes per day (4.6%±1.3% standard error) and the lowest for drinks per week (2.4±0.8%).

**Table 2.**
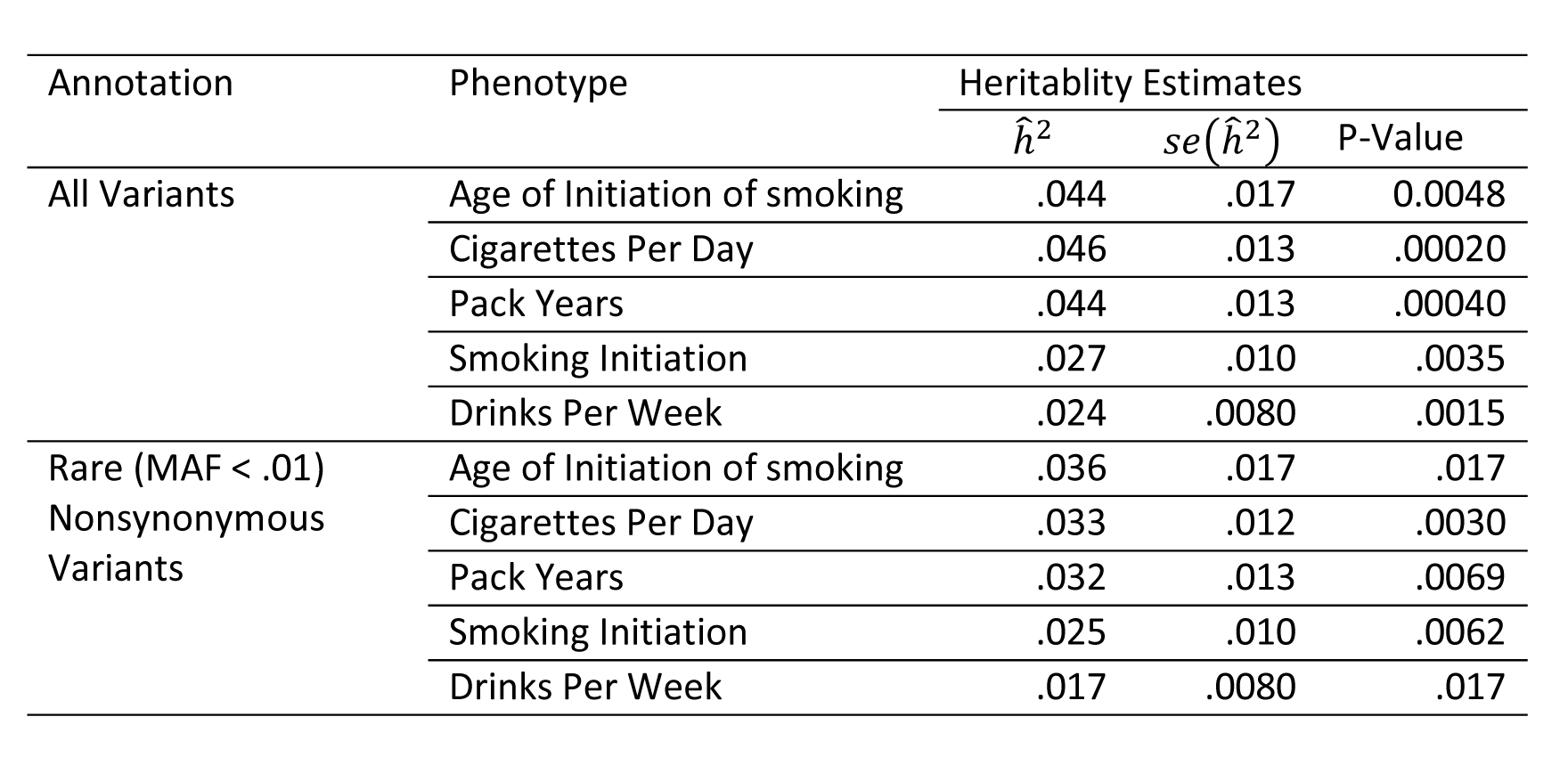
Estimation of Heritablity Explained by Variants on Exome Array. We estimate the heritability based upon a baseline model with 16 different functional categories. The reported heritability ĥ^2^ is based upon the cumulative value from the functional categories with significant heritabilities. We also report the its standard devation (se(ĥ^2^) and p-values, estimated using jackknife.

All pairs of traits are genetically correlated (Table 3) except for cigarettes per day and smoking initiation, and the direction of the genetic correlations are in the expected direction. For instance, cigarettes per day has a positive genetic correlation with drinks per week (0.04±0.0084), consistent with the observation that the increased alcohol consumption is correlated with increased tobacco consumption. Age of initiation has a negative correlation with all other traits, which is consistent with the observation that an earlier age of smoking initation is correlated with increased tobacco and alcohol consumption in adulthood. The patterns and magnitudes of correlation are highly similar when considering only rare nonsynonymous variants (Table 3).

**Table 3.** Estimation of Genetic Correlation Between Smoking and Drinking Traits. We estimate genetic correlations between 5 smoking and drinking traits. Genetic correlation estimates 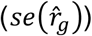, their standard deviation 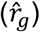 and p-values are reported.

| Trait 1 | Trait 2 | Genetic Correlation |  |  |
| --- | --- | --- | --- | --- |
| | | $\hat{r}_g$ | $se(\hat{r}_g)$ | P-value |
| A. Aggregated Genetic Correlation Induced by All Variants on the Exome Array |  |  |  |  |
| Age of Initiation of Smoking | Cigarettes Per Day | -0.024 | 0.010 | 0.020 |
| Age of Initiation of Smoking | Smoking Initiation | -0.037 | 0.012 | 0.0017 |
| Age of Initiation of Smoking | Drinks Per Week | -0.023 | 0.010 | 0.023 |
| Age of Initiation of Smoking | Pack Years | -0.03 | 0.010 | 0.0040 |
| Cigarettes Per Day | Smoking Initiation | 0.0027 | 0.0088 | 0.76 |
| Cigarettes Per Day | Drinks Per Weeks | 0.040 | 0.0084 | $1.6 \times 10^{-6}$ |
| Cigarettes Per Day | Pack Years | 0.054 | 0.011 | $1.4 \times 10^{-6}$ |
| Smoking Initiation | Drinks Per Week | 0.041 | 0.0058 | $9.4 \times 10^{-13}$ |
| Smoking Initiation | Pack Years | 0.018 | 0.0057 | 0.0012 |
| Drinks Per Week | Pack Year | 0.025 | 0.0070 | 0.00038 |
| B. Genetic Correlation Induced by Rare (MAF < 1%) Nonsynonymous Variants |  |  |  |  |
| Age of Initiation of Smoking | Cigarettes Per Day | -0.024 | 0.010 | 0.020 |
| Age of Initiation of Smoking | Smoking Initiation | -0.033 | 0.011 | 0.0026 |
| Age of Initiation of Smoking | Drinks Per Week | -0.023 | 0.0094 | 0.013 |
| Age of Initiation of Smoking | Pack Years | -0.032 | 0.0088 | 0.00021 |
| Cigarettes Per Day | Smoking Initiation | 0.0025 | 0.0084 | 0.76 |
| Cigarettes Per Day | Drinks Per Weeks | 0.043 | 0.0076 | $1.10 \times 10^{-8}$ |
| Cigarettes Per Day | Pack Years | 0.059 | 0.010 | $1.50 \times 10^{-8}$ |
| Smoking Initiation | Drinks Per Week | 0.013 | 0.0051 | 0.0084 |
| Smoking Initiation | Pack Years | 0.010 | 0.0044 | 0.019 |
| Drinks Per Week | Pack Year | 0.011 | 0.0056 | 0.049 |

## Discussion

With a maximum sample size ranging from 70,847 to 164,142, the present study is the largest study to date of low-frequency nonsynonymous and LoF variants in smoking and alcohol use. Our meta-analytic study design allowed us to conduct single variant, gene-based burden tests, and exact conditional analyses accounting for common variants on the Illumina exome chip and UK Biobank arrays. Despite these analytical advantages and a large sample size, we were unable to discover robust, novel associations for nonsynonymous or LoF variants. The one novel associated rare variant in *STARD3* did not replicate in two complementary large exome chip meta-analysis consortia.

We discovered a common nonsynonymous SNP, rs1260326, in *GCKR,* associated with drinks per week. The T→C change results in a nonsynonymous (Leu→Pro) and splice region change in the final codon of the 14th exon in *GCKR.* The mutation is predicted to be possibly damaging by PolyPhen-2^52^, although the functional significance of this variant is unknown. Denser genotyping or genotype imputation will verify whether this particular variant has a direct causal relationship to drinks per week, or if the association arises artifactually due to linkage disequilibrium between this variant and other variants in the locus. We replicated one of four previous rare single variant associations and twenty five out of twenty six gene-level associations. Possible explanations include the relatively thin coverage of the exome chip compared to targeted resequencing, phenotypic heterogeneity (previous studies used dependence diagnoses), differences in study population, or overestimation of true effects in the original studies.

We showed that rare nonsynonymous variants on the exome chip explain significant proportions of phenotypic variance. The exome chip was designed to genotype coding variants uncovered in ∼12,000 sequenced exomes. By design, it comprehensively ascertained high confidence rare nonsynonymous, splice, and stop variants within those sequences and only sparsely genotypes other classes of variation, including common variants. The use of the exome chip therefore limited our ability to quantify heritability for these other types of variants, or to conduct enrichment tests. Care should also be taken when interpreting those results for which we had substantial coverage on the exome chip. The estimates should be interpreted as “chip heritability”, which is the proportion of heritability that can be tagged by variants on the chip. Even rare nonsynonymous variants may be in linkage disequilibrium with other nearby variants, and thus the percent variance explained by nonsynonymous variants may not be solely attributed to the genotyped variants. Additional fine mapping and denser genotype data is needed to dissect the contribution of any given variant or class of variants. Nonetheless, our results provide preliminary evidence that nonsynonymous variants contribute substantially to the genetic etiology of smoking and drinking.

The exome chip design is an efficent way to accumulate large samples genotyped with a moderate number of low-frequency exonic variants. The effect size spectrum of low-frequency variants on complex traits is poorly understood and, despite our large sample sizes, it may well be that our meta-analysis was underpowered to detect variants with small effects on smoking and alcohol use behaviors^14^. The maximum sample size for cigarettes per day, for example, was N∼75,000. At this sample size, we had 80% power to detect a variant accounting for >.05% of variance. A small effect, but if the variant in question has MAF=0.1%, it translates to a standardized regression weight of 0.5. That is, for every risk allele an individual carries, their expected phenotype increases by ½ of a standard deviation. Such a variant would be highly consequential for the individuals who carry it, and of considerable scientific interest. The result is similar for SI, where N∼165,000, and we had 80% power to detect an odds ratio >1.14 for a variant with MAF=1%. We had 80% power to detect an odds ratio >1.5 for a variant with MAF=0.1%. The present results indicate there are no rare variants on the exome chip with such effects on smoking and alcohol use in European ancestry individuals.

A similar line of reasoning can be used to put the chip heritability results into context. We found that rare nonsynonymous variants contribute to heritability (e.g., ∼3%) in these traits. Rare disease-associated variants are expected to have larger effects than common variants, in the sense that carrying a rare mutation is expected to have a larger phenotypic impact, if only due to purifying selection for deleterious mutations. However, even if the effect is large in that sense, any rare mutation by definition only affects a small number of individuals. Thus, a rare variant with a large effect accounts for a tiny fraction of variation in any common, complex disorder or trait. In the present study, there were ∼130,000 nonsynonymous variants with MAF<1% (Table S3) and they in aggregate appear to account for substantial variation in the phenotypes. So while the present results provide evidence that rare nonsynonymous variants play a significant role in risk for smoking and alcohol use behavior but that individual rare variants associations remain undetectable even at the sample sizes accumulated here.

## Acknowledgements

Research reported in this article was supported by the National Institute on Drug Abuse and the National Human Genome Research Institute of the National Institutes of Health under award numbers R01DA037904 (SIV), R21DA040177 (DJL), R01HG008983 (DJL), and 5T3DA017637-13 (DMB), as well as funding sources listed in the Supplementary Note. This research has been conducted using the UK Biobank Resource under Application Number 6395.

### Financial Disclosures

There are no conflicts to disclose

